# Genome-wide survey of ribosome collision

**DOI:** 10.1101/710491

**Authors:** Peixun Han, Mari Mito, Yuichi Shichino, Satoshi Hashimoto, Tsuyoshi Udagawa, Kenji Kohno, Yuichiro Mishima, Toshifumi Inada, Shintaro Iwasaki

## Abstract

In protein synthesis, ribosome movement is not always smooth and is rather often impeded for numerous reasons. Although the deceleration of the ribosome defines the fates of the mRNAs and synthesizing proteins, fundamental issues remain to be addressed, including where ribosomes pause in mRNAs, what kind of RNA/amino acid sequence causes this pause, and the physiological significance of this slowdown of protein synthesis. Here, we surveyed the positions of ribosome collisions caused by ribosome pausing in humans and zebrafish on a genome-wide level using modified ribosome profiling. The collided ribosomes, *i.e.*, disomes, emerged at various sites: the proline-proline-lysine motif, stop codons, and the 3′ untranslated region (UTR). The number of ribosomes involved in a collision is not limited to two, but rather four to five ribosomes can form a queue of ribosomes. In particular, XBP1, a key modulator of the unfolded protein response, shows striking queues of collided ribosomes and thus acts as a substrate for ribosome-associated quality control (RQC) to avoid the accumulation of undesired proteins in the absence of stress. Our results provide an insight into the causes and consequences of ribosome slowdown by dissecting the specific architecture of ribosomes.

## Introduction

Translation, which serves as the interface between nucleic acids and amino acids in the central dogma of life, is regulated at multiple steps. Among these steps, translation elongation, a process in which ribosomes travel along mRNAs to synthesize peptides by decoding codons, is interrupted by a wide range of factors: downstream secondary RNA structures, amino acid/tRNA availability, and the presence of nascent peptide chains within the ribosome exit tunnel (Buskirk and Green, 2017). The movement of ribosomes across the transcriptome can be directly assessed by a technique called ribosome profiling, in which ribosome-protected mRNA fragments are generated by RNase treatment and assessed by deep sequencing (Ingolia et al., 2009; Brar and Weissman, 2015). Indeed, this approach has revealed the deceleration of ribosomes under a wide range of conditions (Buskirk and Green, 2017).

It has long been speculated that ribosome deceleration causes the queueing of ribosomes. Although ribosome profiling has revealed ribosome queueing under extreme conditions (*e.g.*, upon perturbation of the translation machinery and the depletion of charged tRNAs (Lareau et al., 2014; Young et al., 2015; Woolstenhulme et al., 2015; Schuller et al., 2017; Darnell et al., 2018; Mohammad et al., 2019; Wu et al., 2019)), the queueing of ribosomes has not been observed in naïve cells (Ingolia et al., 2011). It is unclear whether the lack of stacked ribosomes in native conditions originates from the technical limitations of regular ribosome profiling or the possibility of too few ribosomes on an mRNA to cause a ribosome queue.

Ribosome traffic jams should be avoided by cells since the arrested ribosomes could be the source of truncated and deleterious proteins. Indeed, ribosome arrest is under surveillance in cells by the no-go decay (NGD) system for the mRNA clearance and ribosome-associated quality control (RQC) for the nascent polypeptide degradation. Both systems recognize the distinctive architecture of collided ribosomes (or di-ribosomes, disomes) (Simms et al., 2017; Juszkiewicz et al., 2018; Ikeuchi et al., 2019), triggering the ubiquitination of ribosomes by the E3 ligase ZNF598 (yeast Hel2 homolog) and remodeling of the arrested ribosomes, splitting them into two subunits (Matsuo et al., 2017; Sundaramoorthy et al., 2017; Garzia et al., 2017; Juszkiewicz and Hegde, 2017). In the RQC system, the nascent chain is further degraded due to its ubiquitination by the E3 ligase LTN1 (Bengtson and Joazeiro, 2010; Brandman et al., 2012; Defenouillère et al., 2013; Shao et al., 2013) and the addition of a C-terminal alanine-threonine (CAT) tail (CATylation) (Shen et al., 2015; Osuna et al., 2017; Kostova et al., 2017; Sitron and Brandman, 2019). Although the pivotal roles of these surveillance systems have been illustrated by neurodegeneration in mice in the absence of LTN1 (Chu et al., 2009), the endogenous targets of these systems have remained largely unknown.

In this study, we probed ribosome collisions by the genome-wide survey of disomes in humans and zebrafish with modified ribosome profiling. Dissimilar to the results of disome profiling conducted in yeast (Guydosh and Green, 2014; Diament et al., 2018), the clear accumulation of disome footprints at discrete codons showed that unique amino acid motifs and non-optimal context of stop codons are the contributing factors for ribosome deceleration. Furthermore, long ribosome queues (∼5 ribosomes) were observed in vertebrates under normal conditions. Moreover, disome profiling suggests that post-termination ribosomes along the 3′ untranslated region (UTR) could be an obstacle to ribosomes that follow during the elongation process. We found that severe ribosome traffic jams on X-box binding protein 1 (XBP1), a stress response transcription factor (Hetz and Papa, 2018), induce the degradation of undesirable proteins by RQC in the absence of stress. Our approach provides a versatile framework to overview ribosome traffic jams and mRNAs under surveillance in cells.

## Results and discussion

### Disome footprints define ribosome pause sites

To determine the positions of ribosome traffic jams on mRNAs in humans, we modified the ribosome profiling technique. In typical ribosome profiling (hereafter “monosome” profiling), 17-34 nucleotide (nt)-long ribosome fragments generated by nuclease digestion are isolated (Figures 1A and S1A). In the modified method (or “disome” profiling), we focused on collided ribosomes that can be represented by two ribosomes as a unit. As tightly packed ribosome pairs were reported to protect 40-65 nt-long mRNA fragments (Wolin and Walter, 1988; Guydosh and Green, 2014; Subramaniam et al., 2014; Diament et al., 2018; Juszkiewicz et al., 2018; Ikeuchi et al., 2019), the lengths of disome fragments were expected to be in the same range. Thus, we generated deep sequencing libraries from 50-80 nt mRNA fragments after RNase digestion (Figures 1A and S1A). As a reference, monosome profiling libraries were also prepared from the same samples.

**Figure 1.**
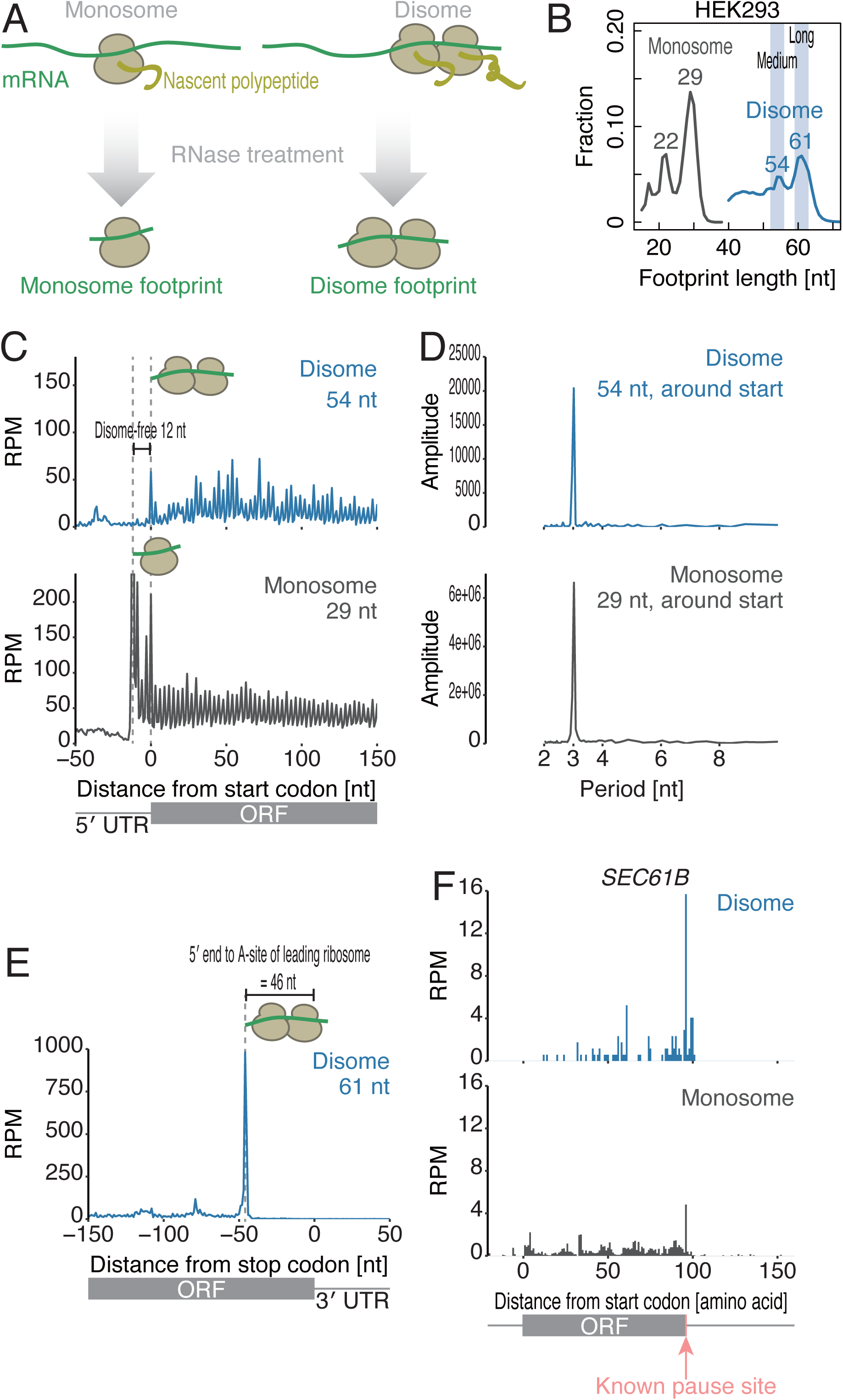
Disome footprints reflect the collision of ribosomes. (A) Schematic representation of disome profiling. (B) Distribution of the lengths of fragments generated by monosome profiling (regular ribosome profiling, gray line) and disome profiling (blue line) in HEK293 cells. (C and E) Metagene analysis showing the 5′ ends of fragments of indicated read lengths around start codons (C) and stop codons (E). The first nucleotide of start or stop codon is set as 0. (D) Discrete Fourier transform of reads showing their periodicity around start codons. (F) The read distribution along *SEC61B* mRNA is depicted, showing the A-site position of a monosome (bottom) and that in the leading ribosome of a disome (top). The known ribosome pause site (Mariappan et al., 2010) is highlighted in pink. RPM: reads per million mapped reads

In our pilot experiments in HEK293 cells, a few specific sites derived from 28S rRNA fragments overpopulated the deep sequencing libraries (Figure S1B and S1C, top). These contaminants are typically generated during RNase digestion, which partially cleaves the rRNAs in ribosomes (Ingolia et al., 2012; Miettinen and Björklund, 2015; McGlincy and Ingolia, 2017; Gerashchenko and Gladyshev, 2017), and escaped from the rRNA depletion step with hybridizing oligonucleotides. To maximize the disome footprint sequencing space in the experiments, we optimized the protocol by introducing Cas9 treatment to degrade specific sequences in the libraries [*i.e.*, Depletion of Abundant Sequences by Hybridization or DASH (Gu et al., 2016)]. Cas9 programed with designed guide RNAs (gRNAs) (Figure S1D) removed the contaminating 28S rRNA fragments from the libraries (Figure S1C, bottom), enabling a prominent increase in the number of usable reads for downstream analysis (Figure S1E). Similar improvement following Cas9 treatment was also observed in monosome profiling (Figure S1F), in which fragments of two different lengths (22 and 29 nt at their peaks) reflecting A-site tRNA accommodation were observed (Figure 1B, gray line) (Wu et al., 2019).

Disome footprints showed the hallmarks of ribosomes in the elongation process. Two populations of mRNA fragments protected by disomes were found to be 54 and 61 nt in length (Figure 1B, blue line), as expected if two consecutive monosome footprints were joined from tail to head. Similar to observations from monosome profiling, discrete Fourier transform showed a 3-nt periodicity in the disome footprints across the coding sequences (CDS) (Figure 1C and 1D), which reflects the codon-wise movement of ribosomes.

Unlike general monosome footprints (Figure 1C, bottom), the disome footprints showed no clear accumulation of the first 12 nt surrounding the start codons by metagene analysis (Figure 1C, top). As discussed in an earlier study (Guydosh and Green, 2014), the scanning ribosome (or initiating 80S ribosome) at the start codon may require this additional region of mRNA; thus, the formation of these fragments would be blocked until the downstream ribosome proceeds forward (Figure S2A). Indeed, the footprints of scanning pre-initiation complexes contain longer fragments than those of the elongating ribosomes (Archer et al., 2016). Similar disome-free regions were drawn from previous studies in yeast, although the disome-free area in yeast was longer (∼25 nt) (Guydosh and Green, 2014). This inconsistency in the length of the disome-free area may be due to the structural differences in ribosomes between mammals and yeast (Anger et al., 2013).

In contrast to the 54 nt disome fragments, which were distributed across the CDS (Figure 1C), the longer 61 nt disome fragments were predominantly found around stop codons (Figure 1E) (note the difference of vertical axis scale with Figure 1C, top), with the 5′ end of the fragments located 46 nt upstream of stop codons. These longer footprints were also found at disomes pause sites (see below for details). These observations represent a scenario in which a leading ribosome pauses at the stop codon and the following ribosome collides into it. Thus, the distance from the 5′ ends to stop codons (46 nt) corresponds to the A-site position of the leading ribosome in the disome and matches the size of one ribosome (∼30 nt) plus the distance from the A-site to the 5′ ends (15 nt) observed in monosome profiling. Similar A-site distance were observed in 54 nt footprints (Figure S3D). This distance is valuable for addressing the A-site positions of the leading ribosomes at subcodon resolution in disome profiling. Indeed, the higher accumulation of disome footprints were found in previously reported translational pause sites at the stop codon of *SEC61B* mRNA, which encodes a tail-anchored (TA) protein (Mariappan et al., 2010; Ingolia et al., 2011) (Figure 1F).

Taken together, these data suggest disome profiling as a platform to survey the landscape of collided ribosomes.

### Widespread disome pause sites across the transcriptome

From disome footprints of both two populations, we identified pronounced pause sites with collided ribosomes (see Experimental Procedures for details) by calculating the disome occupancy at the given codons on which the A-sites of the leading ribosome lay. Here, we defined nearly a thousand sites of ribosome collision (908 disome pause sites in 698 genes) (Figure 2A and Table S1). Given the number of genes (10387) surveyed in the analysis, we estimated that ∼7% of genes have at least one ribosome traffic jam. Although the majority of these traffic jams fell onto the bodies of the mRNA, significant enrichment of ribosome traffic jams at stop codons was observed (Figure 2B, *P* = 1.5 × 10^−177^, hypergeometric test). The distinct disome footprints were exemplified in the representative mRNAs *PRRC2B* and *TXNRD1* (Figure 2C and 2D); strikingly, disome profiling enabled the enrichment of the pausing ribosome fraction and depletion of the ribosome fraction under typical elongation conditions, which dominate monosome profiling. Conversely, regular monosome profiling may miss a fraction of stalled ribosomes, probably because the collided ribosomes were relatively resistant to RNase and not digested into monosomes as previously predicted (Diament et al., 2018).

**Figure 2.**
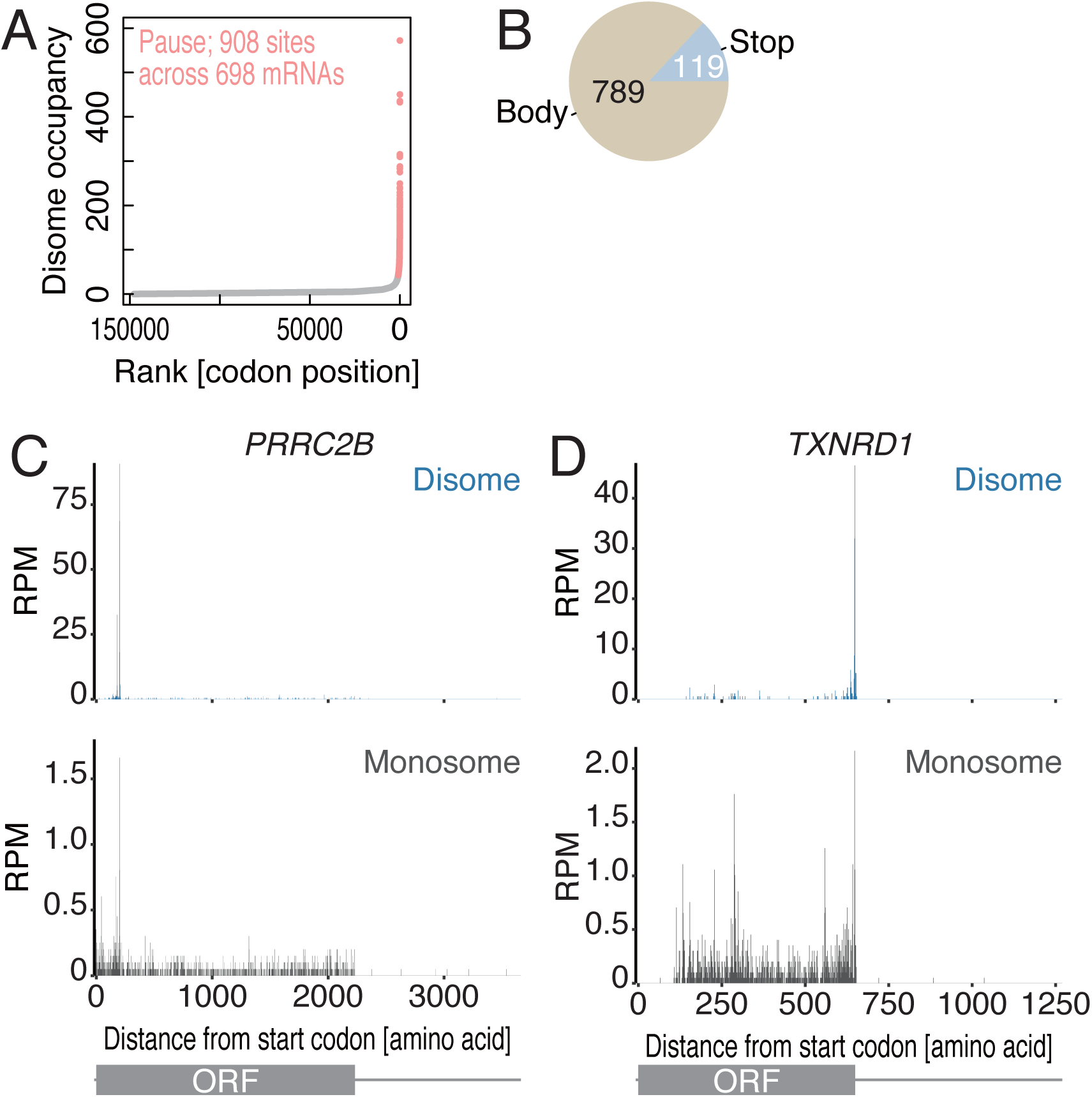
Ribosome collision sites are found in a wide range of mRNAs. (A) Calculated disome occupancies across codons. The collision sites, which are defined as codons with disome occupancies larger than the mean + 3 s.d., are highlighted in pink. (B) Pie chart indicating the positions of disome pause sites in mRNAs. (C and D) The read distributions along *PRRC2B* and *TXNRD1* mRNAs are depicted, showing the A-site position of the monosome (bottom) and that in the leading ribosome of the disome (top).

Next, we investigated the functional implication of the genes associated with ribosome traffic jams and observed that genes encoding chromatin modification factors were much more likely to have disomes (Figure S2B). In contrast, ribosomal genes were not regulated by elongation deceleration; rather, their translation was smooth without ribosome traffic jams.

### Disome pause sites have long ribosome queues

Although disome footprints were generated from two consecutive ribosomes, disome profiling revealed an even longer queue of protein synthesis machinery. Strikingly, the relative disome occupancies around the disome pause sites in the mRNA body and at stop codons showed three upstream peaks every 11 codons (33 nt), which is consistent with the size of a single ribosome (Figure 3A and 3B). Such a long queue of ribosomes was observed for *DDX39B* (Figure S3A) and *XBP1u* mRNAs (Figure 5A). These ribosome queues around disome pause sites were also observed by monosome profiling (Figure S3B and S3C), although they were less clear compared to those in the disome profiles. Ribosome queueing upstream of stop codons was also supported by unbiased metagene analysis around stop codons (Figure S3D), in which discrete Fourier transformed showed a periodicity of 30 nt—the size of a ribosome (Figure S3E). These observations strongly support a model in which the long pause of a ribosome at a specific site causes a long queue of stalled ribosomes even under normal conditions. We noted that the exact number of queued ribosomes could still be under-estimated due to the incomplete RNase digestion of multiple stacked ribosomes, as discussed above.

**Figure 3.**
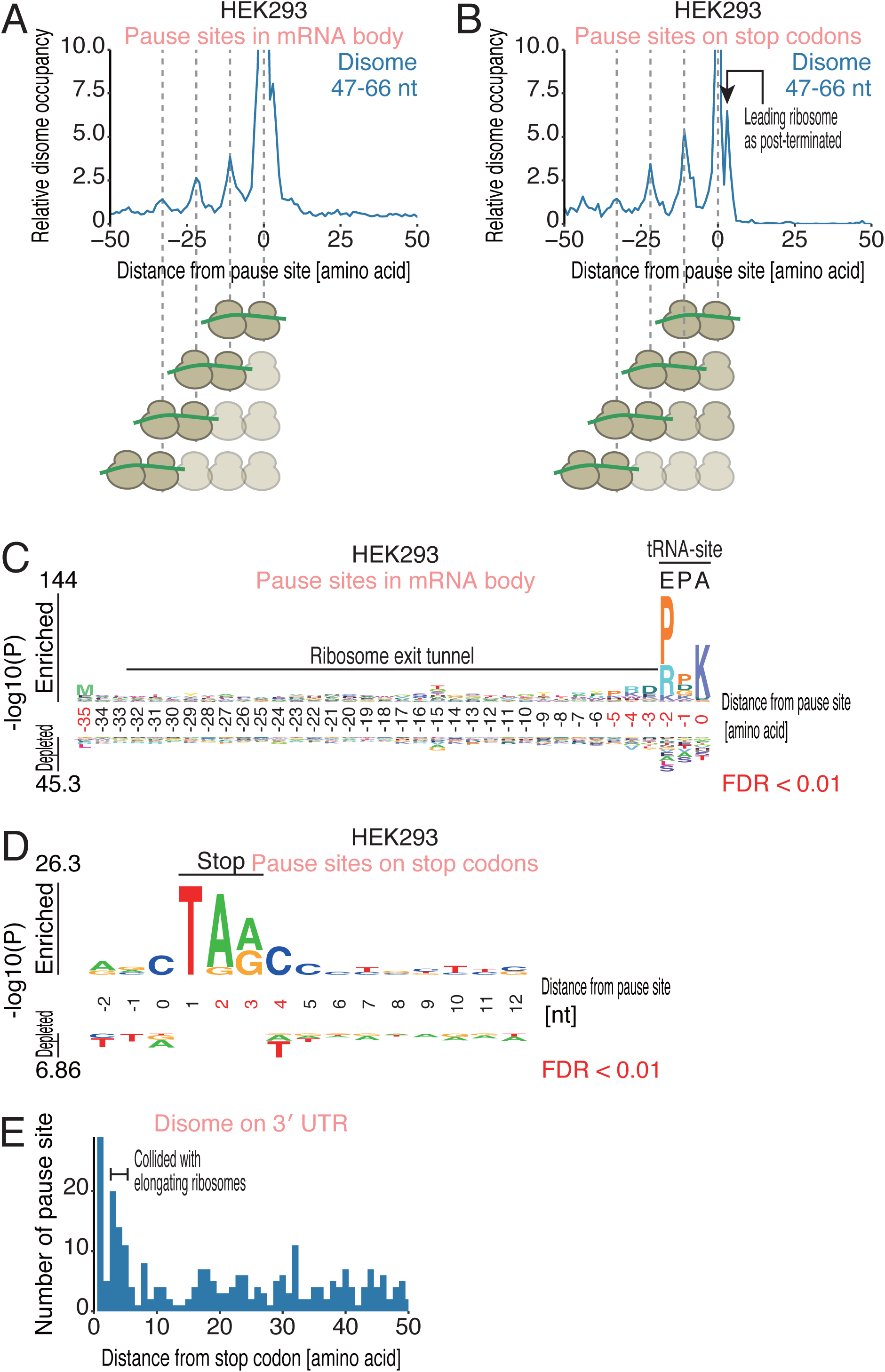
Disome profiling unveils long queues of ribosomes. (A and B) Metagene analysis of disome reads around collision sites in the mRNA body (A) and stop codons (B). The A-site position of the leading ribosome is shown. (C) Amino acid sequences enriched in ribosome collision sites found in the mRNA body. Amino acid relative to the A-site of leading ribosome is shown. (D) Nucleic acid enrichment around ribosome collision sites on stop codons. Nucleic acids from the stop codon (1 as the first nucleotide of stop codon) are depicted. (E) Histogram of the number of disome pause sites found in the 3′ UTR along with the distance from the stop codon. Logos in C and D were drawn by kpLogo (Wu and Bartel, 2017). The positions with significantly enriched sequences are highlighted in red.

**Figure 5.**
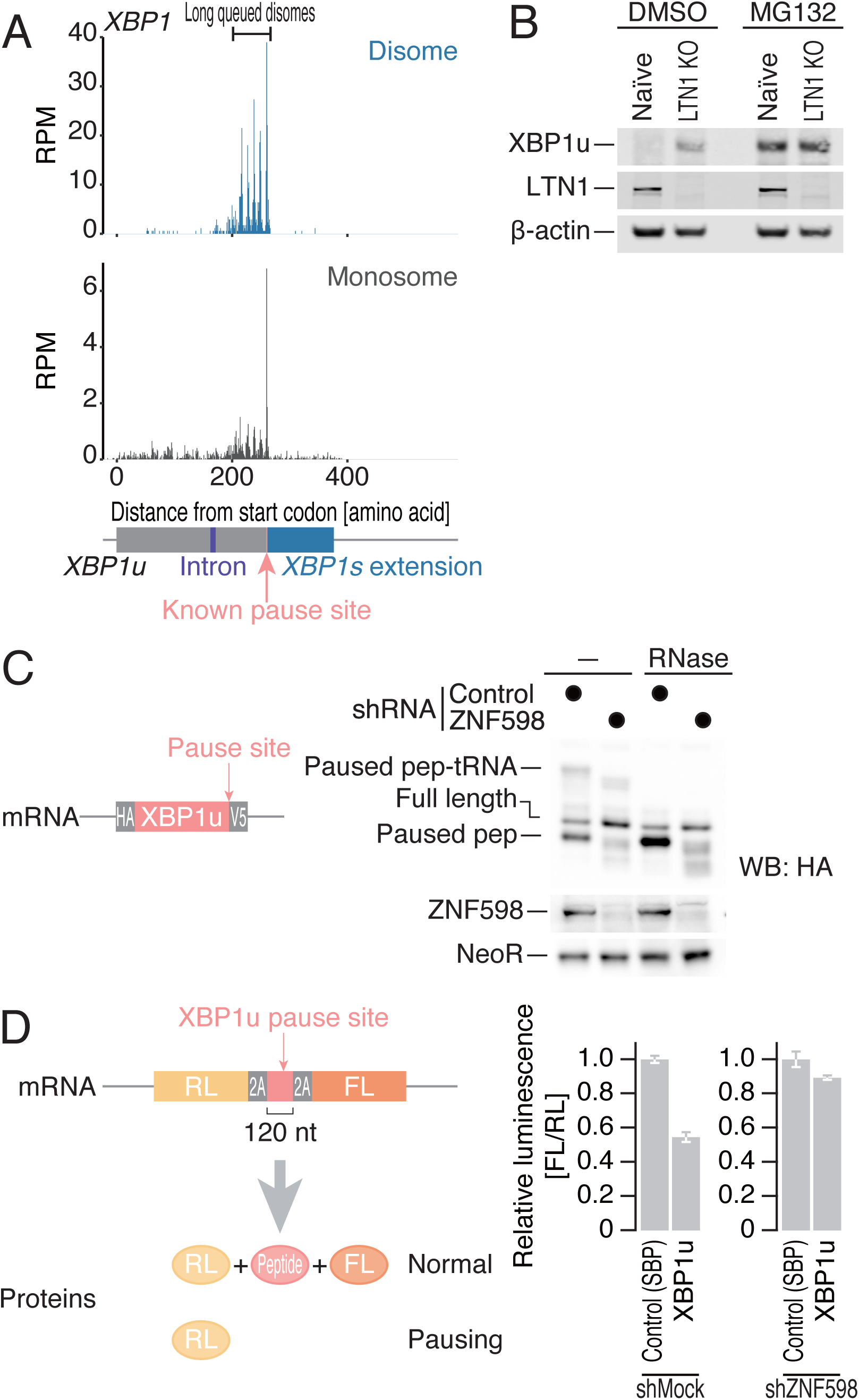
Collided ribosomes on XBP1u are rescued by the RQC pathway. (A) The read distribution along human *XBP1u* mRNA is depicted, showing the A-site position of the monosome (bottom) and that in the leading ribosome of the disome (top). The known ribosome pause site (Yanagitani et al., 2011) is highlighted in pink. (B) Western blot for the indicated proteins in naïve or LTN1 knockout HEK293 cells. (C) Western blot for proteins expressed from a reporter plasmid (left) in ZNF598 knockdown and control cells. Proteins with or without RNase treatment were separated in neutral-pH PAGE to detect the peptidyl-tRNAs. (D) Luciferase assay with the reporter with an XBP1u pause site sandwiched between self-cleavage 2A tags (left) in ZNF598 knockdown and control cells.

### Motifs associated with ribosome collisions

In theory, the high rate of translation initiation—the general limiting step among the overall translation process (Morisaki et al., 2016; Wu et al., 2016; Yan et al., 2016; Wang et al., 2016)—increases the number of ribosomes on mRNAs and thus may lead to ribosome collision. However, ribosome collision was not correlated with translation efficiency (Figure S3F), which was calculated by the over/under-representation of the number of monosome footprints over RNA sequencing (RNA-seq) reads and is generally considered the translation initiation rate (Weinberg et al., 2016). Thus, additional element(s), which consumes even longer time than translation initiation rate, may also determine ribosome collisions in cells.

The motifs around ribosome collision sites potentially explain the origins of ribosome traffic jams. Ribosome collision sites found in the mRNA body showed a strong proline-proline-lysine tendency at the E/P/A-site of the leading ribosome (Figure 3C). The conformation of the nascent chain with a stretch of proline residues is incompatible with peptidyl transferase center (PTC) (Melnikov et al., 2016; Huter et al., 2017) and thus may cause ribosome pausing.

Ribosome collisions at stop codons showed no clear enrichments in nascent chain motif (Figure S4A). Instead, we observed a trend in nucleotides tilted toward a C after the stop codon (+4 position) (Figure 3D). This motif could be attributed to the mechanism of stop codon recognition by eRF1; the +4 purine nucleotide adopts a preferential conformation induced by eRF1 as it interacts with base G626 of 18S rRNA (Brown et al., 2015) and is thus frequently found downstream of open reading frames (ORFs) in eukaryote (Brown et al., 1990). Thus, inefficient translation termination with a C pyrimidine at the +4 site (McCaughan et al., 1995) would cause the collision of upstream translating ribosomes into paused ribosome at the stop codon that had not swiftly dissociated.

Given that the conformational dynamics of ribosomes alter the accessibility of RNase and generate ribosome footprints containing fragments of different sizes (Archer et al., 2016; Mohammad et al., 2016; Wu et al., 2019), we hypothesized that disomes on the stall sites have footprints of characteristic sizes. Among the two populations of fragments in the disome footprints, the 61 nt fragments were found at disome pause sites and queued disomes (Figure S4B-D). Future biochemical and structural dissection of the differences between 54 nt and 61 nt disome fragments is warranted.

### Disomes with post-termination ribosomes

During our analysis of disome accumulation around stop codons, we found a population of disomes spanning the CDS and 3′ UTR (Figure 3B and S3D). Since the A-sites of leading ribosomes lay outside of the CDS, we argued that those disomes represent post-termination ribosomes. Indeed, the disome footprints originated from 3′ UTR did not show 3 nt periodicity, ruling out that disomes in 3′ UTR are under active elongation (Figure S4E). Here, we expanded our survey of disome pause sites along the 3′ UTR and found the enrichment of disome footprint fragments 3-5 amino acids (9-15 nt) downstream of stop codons (Figure 3E). Given the accumulation of post-terminated ribosomes at stop codons and along the 3′ UTR in regular monosome profiling (Guydosh and Green, 2014; Young et al., 2015; Mills et al., 2016; Guydosh and Green, 2017; Sudmant et al., 2018), our data suggested that ribosomes that fail to dissociate after termination and leak into the 3′ UTR could be road blocks for upcoming ribosomes.

### Conservation of ribosome traffic jams in vertebrates

To understand the basis of ribosome collisions across vertebrates, we conducted disome profiling in zebrafish embryos. Identical to the observations made in human cells, zebrafish disome footprint fragments were double the size of a monosome footprint fragments and exhibited a 3-nt periodicity (Figure S5A-C); the first 12 nt of the CDS were disome free (Figure S5B); a 44 nt-long stretch to the A-site of the leading ribosome were determined by reads upstream of stop codons (Figure S5D). The zebrafish disomes also indicated ribosome pausing at the known pause site of *Sec61b* (Figure S5E). Furthermore, we called pause sites *de novo* and found them across mRNA bodies and on stop codons in a distribution similar to that in HEK293 cells (Figure S5F and S5G).

Similar to the results of human disome profiling (Figure 3A), long queues of ribosomes formed upstream of disome pause sites in the zebrafish embryo disome profile (Figure 4A). The identical amino acids enriched at the E/P/A-site of the disome pause sites suggests a common basis for ribosome deceleration in vertebrates (Figure 4B). Although a small number of orthologs between the disome pause sites from the two species were found (Figure 4C, top), the shared orthologs exhibited ribosome collisions at the exact same sites (Figure 4C, bottom), which was exemplified by *HDAC2/hdac1* and *BRD2/brd2a* (Figure 4D). The conservation of ribosome collision between these two species suggests its pivotal functions during protein synthesis.

**Figure 4.**
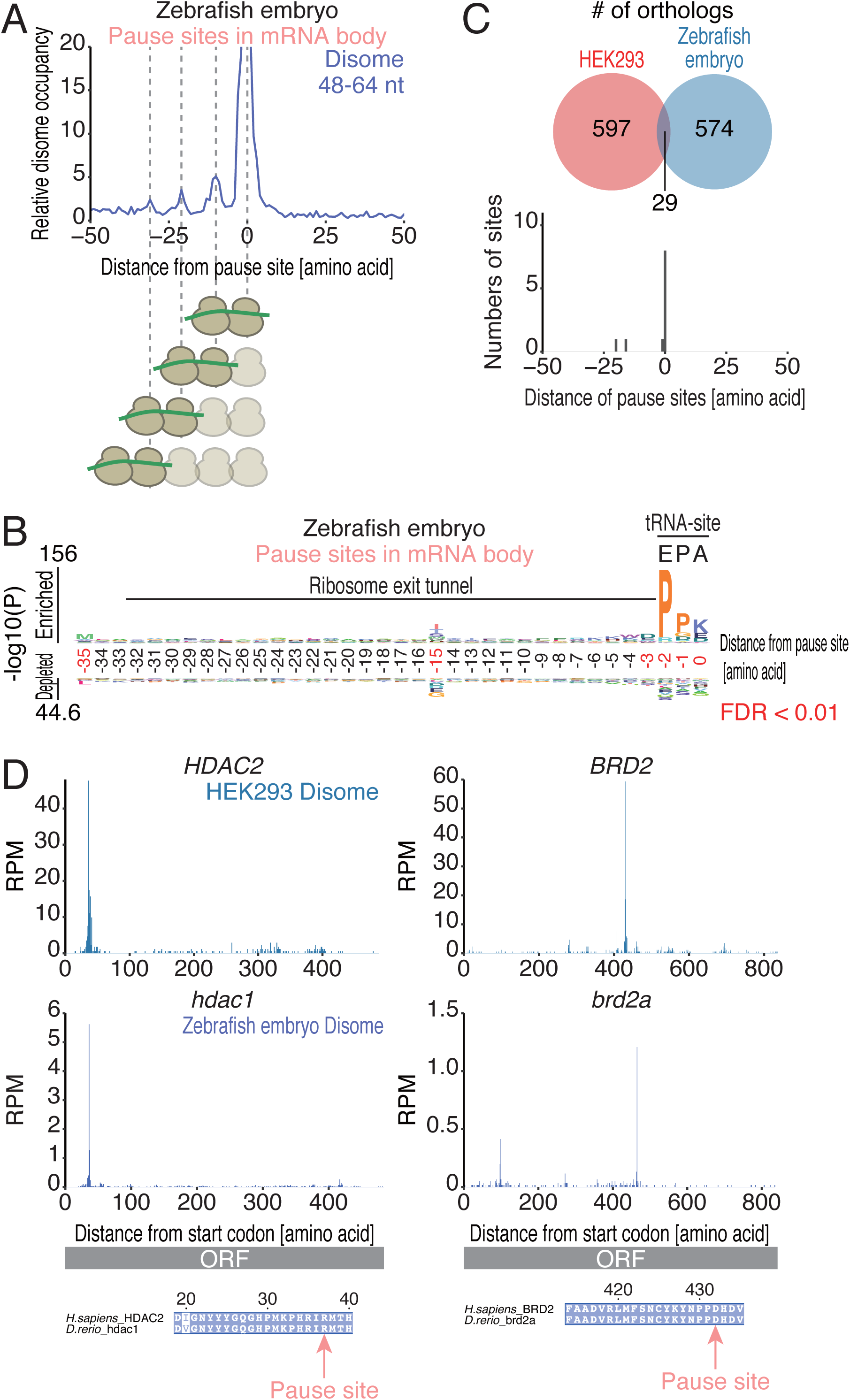
The conservation of ribosome collisions between humans and zebrafish. (A) Metagene analysis of zebrafish disome reads around ribosome collision sites in the mRNA body. (B) Same logo as that in Figure 3C showing zebrafish ribosome collision sites in the mRNA body. (C) Venn diagram showing the overlap of ribosome collisions identified in humans and zebrafish orthologs (top). The distribution of the lengths of collision sites between orthologs is depicted (bottom). (D) Disome footprint distributions in human and zebrafish orthologs [*HDAC2/hdac1* (left) and *BRd2/brd2a* (right)] are depicted, showing the A-site position of the leading ribosome of the disome.

### XBP1u protein was decayed by RQC

We searched for mRNAs that contain significant ribosome queues upstream of disome pause sites. *XBP1u* emerged as the mRNA with the most prominent ribosome queueing features (Figure 5A). We observed five sharp disome peaks (representing 6 ribosomes queued in a row) at 11 amino acid intervals, and the highest peak in the disome footprint occurred at the reported ribosome pause site (Yanagitani et al., 2011) (Figure 5A), at which PTC of ribosome is distorted by the unique conformation of nascent chain within ribosome exit tunnel (Shanmuganathan et al., 2019).

Since ribosome collisions drive RQC (Simms et al., 2017; Juszkiewicz et al., 2018; Ikeuchi et al., 2019), the long queue of ribosome on the *XBP1u* mRNA led us to investigate whether the XBP1u protein is subject to RQC. For this purpose, we knocked out LTN1 (Figure S6A), an E3 ligase that triggers nascent chain ubiquitination and subsequent proteasome degradation (Inada, 2017). Since the XBP1u protein is rapidly degraded through the ubiquitin-proteasome pathway (Lee et al., 2003; Yoshida et al., 2006), no protein accumulation was observed (Figure 5B). In contrast, LTN1 depletion (or the treatment of proteasome inhibitor MG132) rescued the protein from degradation (Figure 5B).

The RQC pathway is initiated by the recruitment of the E3 ligase ZNF598 (yeast Hel2 homolog) to collided ribosomes (Simms et al., 2017; Juszkiewicz et al., 2018; Ikeuchi et al., 2019), which is then followed by the dissociation of the stalled ribosome, leaving paused peptidyl-tRNA in the 60S subunit. When ZNF598 does not recognize the collided ribosomes, the paused ribosomes read through the pause site and translate the downstream regions (Juszkiewicz and Hegde, 2017; Sundaramoorthy et al., 2017; Matsuo et al., 2017; Juszkiewicz et al., 2018). The XBP1u pause site phenocopied the reported fate of the RQC target; ZNF598 knockdown alleviated the accumulation of paused peptidyl-tRNA, which is observed as the higher bands that disappeared with the RNase treatment, and produced more full-length proteins than those observed in control cells (Figure 5C). Moreover, similar downstream translation was also recapitulated with the short pausing sequence inserted into the middle of the N-terminal *Renilla* and C-terminal firefly luciferase ORFs (Figure 5D). We noted that the flanking viral 2A sequences, which induces a self-cleavage of peptide bond during translation (de Felipe et al., 2006; Juszkiewicz and Hegde, 2017; Sundaramoorthy et al., 2017), liberated the individual luciferase proteins and ensured the effect of ribosome stalling in our output, avoiding to reflect protein degradation by RQC (Juszkiewicz and Hegde, 2017; Sundaramoorthy et al., 2017). XBP1u translation is essential to tether the mRNA-ribosome-nascent chain complex onto the endoplasmic reticulum (ER) membrane until the unfolded protein response (UPR) is induced (Yanagitani et al., 2009; Yanagitani et al., 2011; Kanda et al., 2016; Shanmuganathan et al., 2019), which leads to unconventional splicing of *XBP1u* mRNA anchored on ER (Yoshida et al., 2001; Calfon et al., 2002) and a frame shift to produce XBP1s protein. However, the protein product of the unspliced form of *XBP1* is undesirable under normal conditions. Thus, cells harness the RQC system to degrade the unwanted transcription factor in the absence of stress. We note that the intron of XBP1u is not covered by stacked ribosomes (Figure 5A), thus allowing its splicing upon endoplasmic reticulum (ER)-stress without steric hindrance by ribosomes.

### To pause or not to pause

Ribosome stacking is not rigid but is rather flexible, probably due to diverse factors in cells, such as the kinetics of translation initiation, elongation, termination, and recycling, which finally determine the number of ribosomes in a queue. Indeed, in zebrafish embryos, we observed the buildup of disomes at the *xbp1u* pause site (Figure S7A) and the inhibition of downstream translation by the pausing sequence on reporter mRNAs (Figure S7B-D). However, the queueing of ribosomes upstream of the *xbp1u* pause site was not obvious (Figure S7A). Therefore, we would not be surprised if the number of queued ribosomes is different, depending on cell type and species.

The sorting of ribosomes for RQC and ribosome queueing should be related but distinct issues. Whereas we found that the ribosome collision on *XBP1u* is rescued by RQC, ribosome collisions was not enough to predict the RQC substrate; a readthrough of ribosome collision sites in ZNF598 knockdown cells was not observed in monosome profiling (Figure S6B), suggesting that the substrate selection in RQC requires extra determinants.

In addition to their role in quality control, ribosome pauses are suggested to help cotranslational nascent protein folding (Komar, 2009). Thus, ribosome collision could be a double-edged sword for protein synthesis and decay. How ribosome pauses in specific genes (with GO terms shown in Figure S2B) affect cellular outcomes should be studied in the future. The strikingly versatile disome profiling will expand our knowledge regarding the *in vivo* kinetics of elongation and termination, which have been underestimated by regular monosome profiling, and the fates of mRNAs and proteins defined by cotranslational events.

## Acknowledgments

We thank all the members of the Iwasaki laboratory for constructive discussion, technical help, and critical reading of the manuscript. S.I. was supported by a Grant-in-Aid for Scientific Research on Innovative Areas “nascent chain biology” (JP17H05679) and a Grant-in-Aid for Young Scientists (A) (JP17H04998) from JSPS, the Pioneering Projects (“Cellular Evolution”) and the Aging Project from RIKEN, and the Takeda Science Foundation. YM was supported by a Grant-in-Aid for Scientific Research on Innovative Areas “nascent chain biology” (JP17H05662) from MEXT and Grant-in-Aid for Scientific Research (B) (JP18H02370) from JSPS. T.I. was supported by a Grant-in-Aid for Scientific Research (KAKENHI) from the Japan Society for the Promotion of Science (Grant number 18H03977 to TI) and by the Takeda Science Foundation. DNA libraries were sequenced by the Vincent J. Coates Genomics Sequencing Laboratory at UC Berkeley, supported by an NIH S10 OD018174 Instrumentation Grant. Computations were supported by Manabu Ishii, Itoshi Nikaido, and the Bioinformatics Analysis Environment Service on RIKEN Cloud at RIKEN ACCC. YS was a JSPS Research Fellow (PD) (19J00920).

## Author contributions

P.H., Y.S., M.M., and S.I. performed experiments; P.H. and Y.S. analyzed the deep sequencing data; S.H. and T.I. established the cell lines used in this study and analyzed peptidyl-tRNA conjugate with the input from K.K; T.U. and T.I. performed monosome profiling with ZNF598 knock-down; zebrafish disome profiling was helped by Y.M; S.I. supervised the project; P.H. and S.I. wrote the manuscript with the edit from all the authors.

## Experimental procedures

### Cell lines

HEK293 cells were cultured in DMEM + GlutaMAX-I (Thermo Fisher Scientific) with 10% FBS at 5% CO_2_ at 37°C. Cells were treated with 0.5 µM MG132 (Wako Chemicals) dissolved in DMSO for 6 hr.

LTN1 knockout cells were generated using CRISPR-Cas9 approaches (Ran et al., 2013). The guide sequence used to target LTN1 locus was 5′- ATTCCACCACAACCTAACCATGG-3′ (Figure S6A) and was cloned into the BbsI restriction sites in the pSpCas9(BB)-2A-Puro (PX459) plasmid. HEK293T cells were transfected with the plasmid and selected with 10 µg/ml puromycin, then single cell clone was isolated by limiting dilution and screened by LTN1 expression by western blotting. ZNF598 knockdown cells were generated using shRNA expressing lentivirus (Matsuo et al., 2017). The shRNA against ZNF598 (designed to express 5′- GCCAGTTGCCGTCGTCGTTAAT-3′ guide RNA (Matsuo et al., 2017)) was cloned into HpaI/XhoI restriction sites of pll3.7 plasmid whose GFP gene was substituted to puromycin resistant gene. HEK293T cells infected with shZNF598-expressing lentivirus were selected with 10 µg/ml puromycin.

### Monosome and disome profiling

#### Library preparations

HEK293 cell disome and monosome profiling libraries were prepared as described previously (McGlincy and Ingolia, 2017) with some modifications. Protected RNA fragments ranging from 17-34 nt and 50-80 nt were gel-excised for monosome and disome profiling, respectively.

Zebrafish embryos were obtained by natural bleeding of the AB strain. For the ribosome footprint analysis, 50-60 embryos were snap-frozen at the sphere stage by liquid nitrogen and with lysed with lysis buffer (McGlincy and Ingolia, 2017). These embryos were injected with mRNA libraries of GFP reporters containing various codon-tag sequences. Detailed information about the library and related results will be described elsewhere (Mishima and Iwasaki, in preparation). To isolate ribosomes, an S-400 HR gel filtration spin column (GE Healthcare) was used instead of sucrose cushion ultracentrifugation. RNA fragments ranging from 26-34 nt and 50-80 nt were selected for monosome, and disome profiling, respectively.

PCR-amplified DNA libraries were further treated with gRNA-programmed Cas9 protein. To prepare gRNAs by *in vitro* transcription, template DNAs were amplified by PCR (PrimeSTAR Max, Takara) with the 4 overlapping DNA oligos (three scaffolding oligos and one target-specific oligo) listed below:

scaffold oligo DNAs:

gRNA scaffold 1 (T7 promoter): 5′-TGACTAATACGACTCACTATAGG-3′, gRNA scaffold 2: 5′-

AAAAAAAGCACCGACTCGGTGCCACTTTTTCAAGTTGATAACGGACTAGCCTTATTTAAACTTGCTATGCTGTTTCCAGC-3′, and

gRNA scaffold 3: 5′-AAAAAAAGCACCGACTCGGTGC-3′;

target-specific oligo DNAs:

Disome g1: 5′- TAATACGACTCACTATAGGCCGGTCGCGGCGCACCGCCGGTTTAAGAGCTA TGCTGGAAACAGCATAGCAAGTTTAAATAAGG-3′,

Disome g2: 5′- TAATACGACTCACTATAGGTCGCCGAATCCCGGGGCCGAGTTTAAGAGCTA TGCTGGAAACAGCATAGCAAGTTTAAATAAGG-3′,

Disome g3: 5′- TAATACGACTCACTATAGGAGGCCTCTCCAGTCCGCCGAGTTTAAGAGCTAT GCTGGAAACAGCATAGCAAGTTTAAATAAGG-3′,

Monosome g1: 5′- TGACTAATACGACTCACTATAGGCGGAGGATTCAACCCGGCGGGTTTAAGA GCTATGCTGGAAACAGCATAGCAAGTTTAAATAAGG-3′,

Monosome g2: 5′- TGACTAATACGACTCACTATAGGACGCCGGCGCGCCCCCGCGGTTTAAGAG CTATGCTGGAAACAGCATAGCAAGTTTAAATAAGG-3′,

Monosome g3: 5′- tgacTAATACGACTCACTATAGGGCCGGGCCACCCCTCCCAGTTTAAGAGCTA TGCTGGAAACAGCATAGCAAGTTTAAATAAGG-3′,

Monosome g4: 5′- TGACTAATACGACTCACTATAGGCCCGGGCGGGTCGCGCCGTCGTTTAAGA GCTATGCTGGAAACAGCATAGCAAGTTTAAATAAGG-3′,

Monosome g5: 5′- TGACTAATACGACTCACTATAGGCCGGCCGAGGTGGGATCCCGGTTTAAGA GCTATGCTGGAAACAGCATAGCAAGTTTAAATAAGG-3′,

Monosome g6: 5′- TGACTAATACGACTCACTATAGGACGGGCCGGTGGTGCGCCCTGTTTAAGAGCTATGCTGGAAACAGCATAGCAAGTTTAAATAAGG-3′,

Monosome g7: 5′- TGACTAATACGACTCACTATAGGCGCTTCTGGCGCCAAGCGCCGTTTAAGAGCTATGCTGGAAACAGCATAGCAAGTTTAAATAAGG-3′,

Monosome g8: 5′- TGACTAATACGACTCACTATAGGTTGGTGACTCTAGATAACCTGTTTAAGAGCTATGCTGGAAACAGCATAGCAAGTTTAAATAAGG-3′,

Monosome g9: 5′- TGACTAATACGACTCACTATAGGCGCTCAGACAGGCGTAGCCCGTTTAAGAGCTATGCTGGAAACAGCATAGCAAGTTTAAATAAGG-3′, and

Monosome g10: 5′- TAATACGACTCACTATAGGGCCCAAGTCCTTCTGATCGGTTTAAGAGCTATGCTGGAAACAGCATAGCAAGTTTAAATAAGG-3′.

Purified DNAs were used for *in vitro* transcription with a T7-Scribe Standard RNA IVT Kit (CELLSCRIPT). To generate the disome profiling library, an RNA-protein complex was formed with 1 µM Cas9 protein (New England Biolabs) and 1 µM gRNA pool (3 gRNAs in total) in 4× Cas9 Reaction Buffer (New England Biolabs) at 25°C for 10 min in a 5 µl reaction volume. The reaction mixture was directly added to 15 µl of DNA library containing 0.17 pmol DNA (thus diluting the Cas9 Reaction Buffer to 1×) and incubated at 37°C for 30 min. To generate the monosome profiling library, 2 µM Cas9 protein and 2 µM gRNA pool (10 gRNAs in total) were preincubated and used to generate 0.28 pmol of DNA library. To quench the reaction, gRNAs were degraded by the addition of 0.5 µl of RNaseI (10 U/µl, Epicentre) and incubation at 37°C for 10 min, and then Cas9 proteins were digested with 0.5 µl of proteinase K (∼20 mg/ml, Roche) at 37°C for 10 min. DNA was further gel-excised and sequenced with the HiSeq4000 platform (Illumina).

#### Data analysis

Data were processed as previously described (Ingolia et al., 2012; Iwasaki et al., 2016). Disome and monosome occupancies were calculated as the ratio of reads at given codons to average reads per codon on the transcript. Gene ontology analysis was performed with iPAGE (Goodarzi et al., 2009). The highest disome occupancy was used as a representative value for the analysis. All custom scripts used in this study are available upon request.

For calculation of the translation efficiencies, the read number of ribosome profiling within each CDS were normalized by the read number of RNA-seq using the DESeq package (Anders and Huber, 2010). Reads corresponding to the first and last five codons of each CDS were omitted.

### DNA constructions

#### psiCHECK2-2A-3xFLAG-XBP1u_pause-2A and 2A-3xFLAG-SBP-2A

DNA fragments with the following sequence were inserted between the *Renilla luciferase* (hRluc) and firefly luciferase (luc+) ORFs in the psiCHECK2-EIF2S3 5′ UTR (Iwasaki et al., 2016) to generate an in-frame ORF containing both luciferase ORFs:

2A-3xFLAG-XBP1u_pause-2A: 5′- GGAAGCGGAGCTACTAACTTCAGCCTGCTGAAGCAGGCTGGAGACGTGGAGGAGAACCCTGGACCTGACTACAAAGACCATGACGGTGATTATAAAGATCATGACATCGATTACAAGGATGACGATGACAAGGCCTGGAGGAGCTCCCAGAGGTCTACCCAGAAGGACCCAGTTCCTTACCAGCCTCCCTTTCTCTGTCAGTGGGGACGTCATCAGCCAAGCTGGAAGCCATTAATGAACTCATTCGTTTTGACCGGAAGCGGAGCTACTAACTTCAGCCTGCTGAAGCAGGCTGGAGACGTGGAGGAGAACCCTGGACCT-3′ and

2A-3xFLAG-SBP-2A: 5′- GGAAGCGGAGCTACTAACTTCAGCCTGCTGAAGCAGGCTGGAGACGTGGAGGAGAACCCTGGACCTGACTACAAAGACCATGACGGTGATTATAAAGATCATGACATCGATTACAAGGATGACGATGACAAGGACGAGAAAACCACCGGCTGGCGGGGAGGCCACGTGGTGGAAGGGCTGGCAGGCGAGCTGGAACAGCTGCGGGCCAGACTGGAACACCACCCCCAGGGCCAGAGAGAGCCTAGCGGCGGAGGAGGAAGCGGAGCTACTAACTTCAGCCTGCTGAAGCAGGCTGGAGACGTGGAGGAGAACCCTGGACCT-3′.

PCR products generated from the plasmids and the primers (5′- TGACTAATACGACTCACTATAGG-3′ and 5′- TGTATCTTATCATGTCTGCTCGAAG-3′) were used as templates for *in vitro* transcription.

#### pcDNA3.1(+)-HA-XBP1u-V5

DNA fragments with following sequence were inserted to pcDNA3.1(+) at NheI-ApaI restriction sites:

5’-gccgccaccATGGGATACCCATACGACGTCCCAGACTACGCG**AAGCTTGGTACCGGATCCGAATTT***gtggtggtggcagccgcgccgaacccggccgacgggacccctaaagttctgcttctg tcggggcagcccgcctccgccgccggagccccggccggccaggccctgccgctcatggtgccagcccagagaggggccagcccggaggcagcgagcggggggctgccccaggcgcgcaagcgacagcgcctcacgcacctgagccccgaggagaaggcgctgaggaggaaactgaaaaacagagtagcagctcagactgccagagatcgaaagaaggctcgaatgagtgagctggaacagcaagtggtagatttagaagaagagaaccaaaaacttttgctagaaaatcagcttttacgagagaaaactcatggccttgtagttgagaaccaggagttaagacagcgcttggggatggatgccctggttgctgaagaggaggcggaagccaaggggaatgaagtgaggccagtggccgggtctgctgagtccgcagcactcagactacgtgcacctctgcagcaggtgcaggcccagttgtcacccctccagaacatctccccatggattctggcggtattgactcttcagattcagagtctgatatcctgttgggcattctggacaacttggacccagtcatgttcttcaaatgcccttccccagagcctgccagcctggaggagctcccagaggtctacccagaaggacccagttccttaccagcctccctttctctgtcagtggggacgtcatcagccaagctggaa gccattaatgaac***GAATTCGATATCACCGGTCTCGAGTCTAGA**GGTAAGCCTAT CCCTAACCCTCTCCTCGGTCTCGATTCTACGTAA-3’, where the lowercase italic indicates XBP1u CDS, uppercase characters indicate HA/V5 tag sequence, and bold letters indicate multicloning site sequence.

#### pCS2+sfGFP-zXbp1opt-suv39h1a 3’ UTR

A DNA fragment encoding sfGFP ORF lacking a stop codon and containing EcoRI-XhoI sites at its 3’ end was synthesized by Thermo Fisher Gene ArtStrings service and inserted at NcoI-XhoI sites of pCS2+EGFP-suv39h1a (Mishima and Tomari, 2016) by replacing EGFP ORF (pCS2+sfGFP-suv39h1a). DNA oligos corresponding to zebrafish *xbp1* C-terminal pause sequence or its mutant, as shown below, were annealed and inserted at XhoI-XbaI sites of pCS2+sfGFP-suv39h1a. The pause sequence was codon-optimized to the zebrafish genome to minimize the effect of synonymous codon choice on mRNA stability (Mishima and Tomari, 2016).

y750zxbppauseOptRIXIwtFW: 5’-aattcCAGGAGAGCAAGTACCTGCCTCCGCACCTGCAGCTGTGGGGCCCGCACCAGCTGAGCTGGAAGCCCCTGATGAACTGAc- 3’,

y750zxbppauseOptRIXIwtRV: 5’-tcgagTCAGTTCATCAGGGGCTTCCAGCTCAGCTGGTGCGGGCCCCACAGCTGCAGGTGCGGAGGCAGGTACTTGCTCTCCTGg-3’,

y751zxbppauseOptRIXImutFW: 5’-aattcCAGGAGAGCAAGTACCTGCCTCCGCACgcaCAGCTGTGGGGCCCGCACCAGCTGAGCgcaAAGCCCCTGATGAACTGAc-3’,

and

y751zxbppauseOptRIXImutRV: 5’-tcgagTCAGTTCATCAGGGGCTTtgcGCTCAGCTGGTGCGGGCCCCACAGCTGtgcGTGCGGAGGCAGGTACTTGCTCTCCTGg-3’

### Reporter assay

Reporter mRNAs were prepared and transfected as previously described (Iwasaki et al., 2016; Iwasaki et al., 2019). Luminescence was detected with a dual-luciferase reporter assay system (Promega) and GloMax (Promega).

### Neutral-PAGE

For neutral-PAGE, cells were lysed by passive lysis buffer (Promega) and centrifuged at 13,500 g for 1 min. Supernatants were collected and same amounts of total proteins were used as protein samples. For RNase (+) samples, RNase A (QIAGEN, 19101) was added at a final concentration of 0.05 µg/µl and incubated on ice for 20 min: for RNase (-) samples, Milli-Q water was added instead. After incubation, NuPAGE sample buffer (200 mM Tris-HCl pH6.8, 8% w/v SDS, 40% glycerol, 0.04% BPB, and 100 mM DTT) was added and heated at 65°C for 10 min. Proteins were separated by 15% PAGE at neutral pH condition (pH 6.8) for 4 hr at 50 mA constant current with MES-SDS buffer (1 M MES, 1 M Tris base, 69.3 mM SDS, and 20.5 mM EDTA) and were transferred to a PVDF membrane (Millipore, IPVH00010).

### Western blot analysis

The anti-XBP1 (Cell Signaling Technology, 12782), anti-LTN1 (Abcam, ab104375), anti-β-actin (Medical & Biological Laboratories, M177-3), anti-HA-Peroxidase (Roche, 12013819001), anti-ZNF598 (Novus, NBP1-84658), anti-neomycin phosphotransferase II (Merck KGaA, 06-747), anti-GFP (MBL, 598), and anti-tublin (Sigma-Aldrich, T6074) primary antibodies were used for Western blotting. To generate the Western blot shown in Figure 6B, IRDye680- or IRDye800-conjugated secondary antibodies (LI-COR, 925-68070/71 and 926-32210/11) were used to detect proteins, and images were collected by an Odyssey CLx Infrared Imaging System (LI-COR). To generate the Western blot shown in Figure 6C, HRP-Linked anti-rabbit IgG antibodies (GE Healthcare, NA934) were used for the detection and chemiluminescence were collected by LAS 4000 mini (GE Healthcare). To detect zebrafish embryo proteins by Western blotting, HRP-conjugated anti-rabbit IgG (MBL, 458) and HRP-conjugated anti-mouse IgG (MBL, 330) were used. Signals were detected by LuminaForte (MerkMillipore) and Amersham Imager (GE Healthcare).

### Zebrafish microinjection

Capped and polyadenylated sfGFP-Xbp1 mRNAs were *in vitro* synthesized as described previously (Mishima and Tomari, 2016) and injected into 1-cell stage embryos at the concentration of 50 ng/µl.

### Accession numbers

The results of monosome profiling, disome profiling, and RNA-seq of HEK293 cells (GSE133393 and GSE126298) and zebrafish (GSE133392) used in this study were deposited in the National Center for Biotechnology Information (NCBI).

---

**Figure S1.**
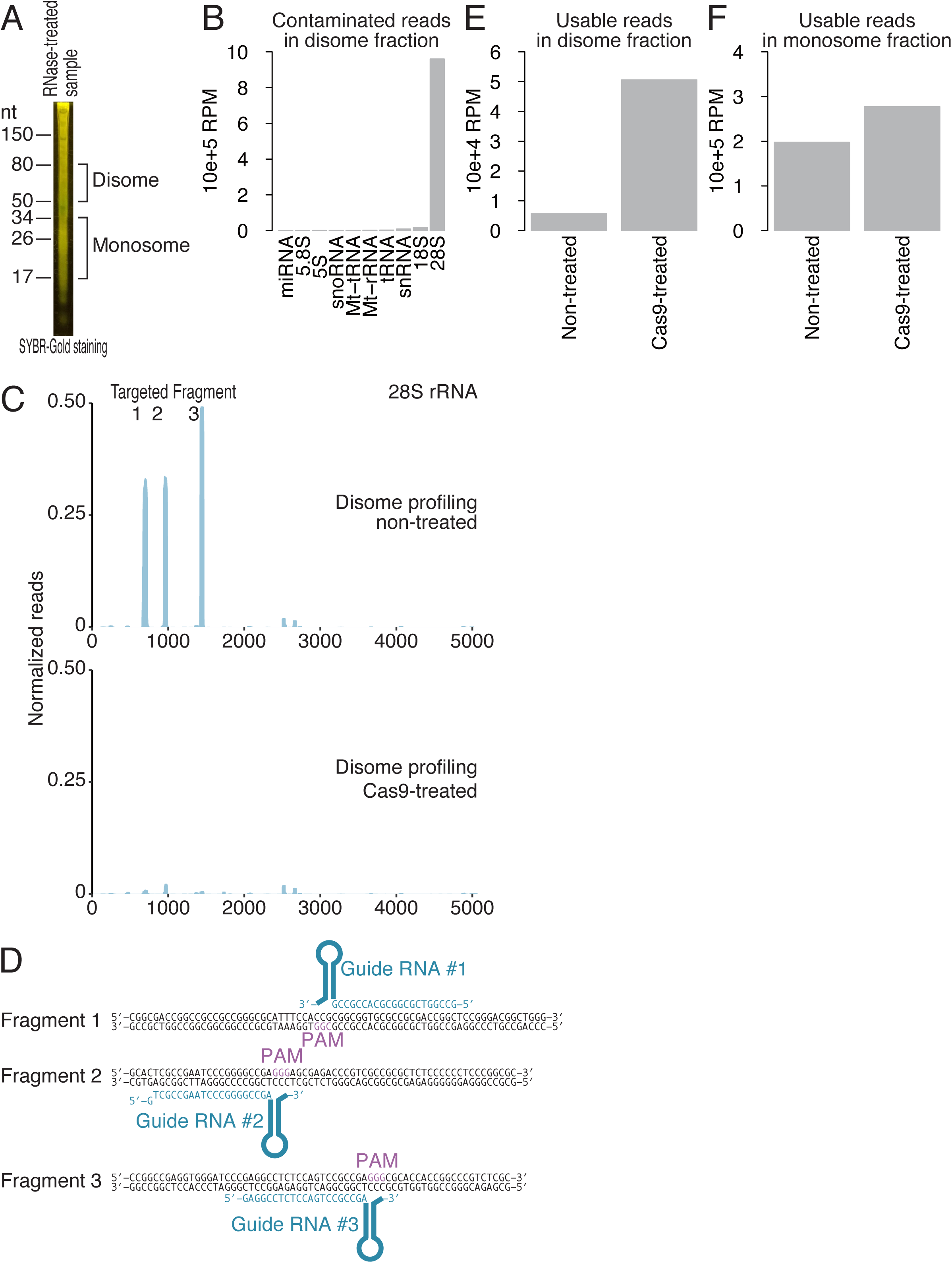
CRISPR-Cas9–mediated depletion of contaminated 28S fragments improves the yield of disome footprints. (A) SYBR-Gold staining of RNAs protected from RNaseI treatment. Regions of RNA gel-excised for monosome and disome profiling are indicated. (B) The origin of the contaminated reads in Cas9-untreated disome profiling libraries. (C) Read distribution of 28S rRNA in untreated (top) and Cas9-treated (bottom) disome profiling libraries. Reads are normalized to those originating from the 2493-2544 nt 28S rRNA region, which is not targeted by gRNAs. (D) gRNAs to eliminate three 28S fragment contaminants dominating the disome libraries. (E) Bar graph showing disome profiling reads that mapped to the genome but not to non-coding RNAs indicated in Figure S1B. Cas9 treatment improved the yields of usable disome footprints, which reflect relatively small fraction of ribosomes in cells, for downstream analysis. (F) Same as (E), but for monosome profiling. gRNAs were designed to eliminate contaminated rRNAs in monosome profiles. RPM: reads per million mapped reads

**Figure S2.**
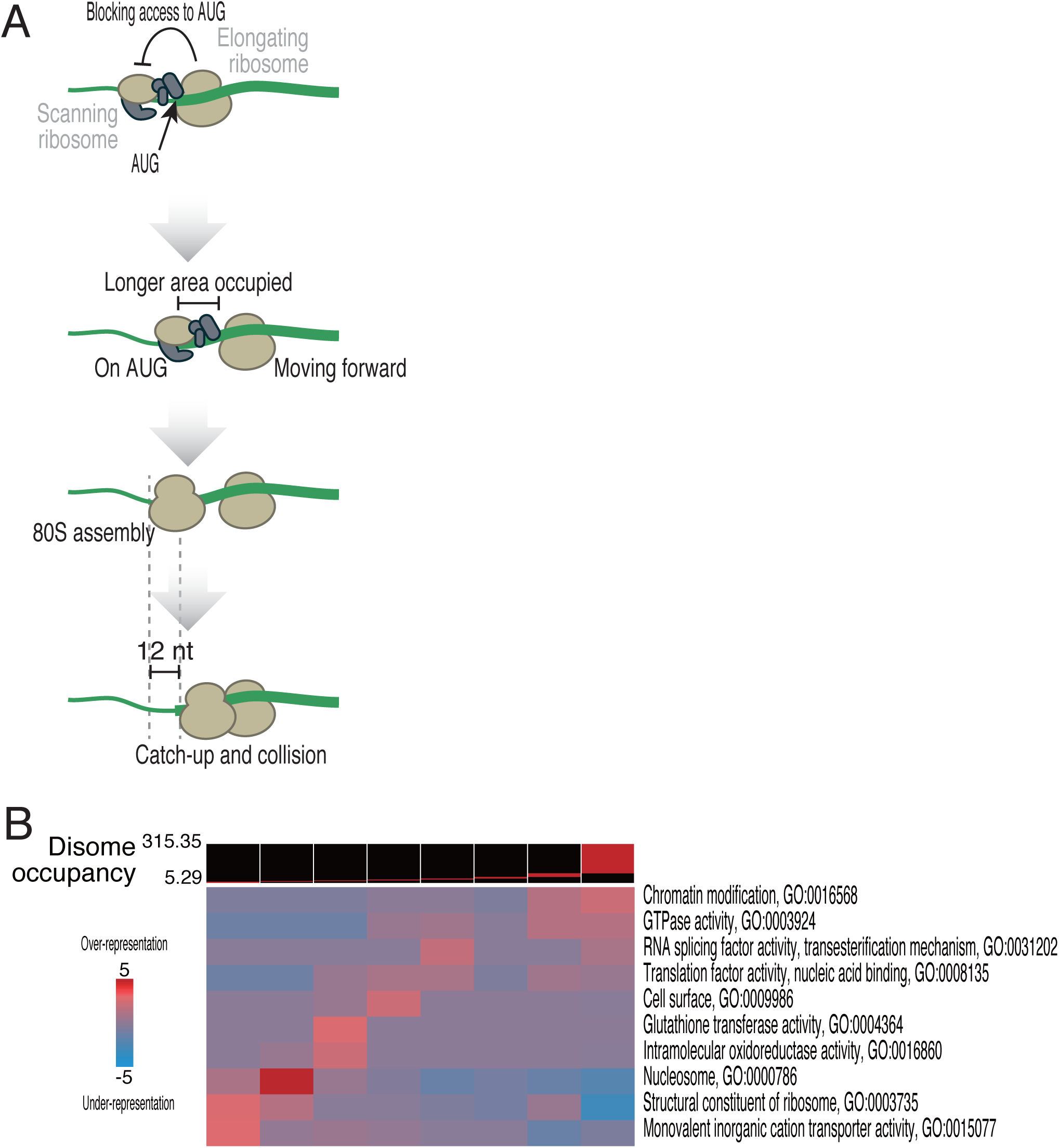
Gene ontology pathways associated with ribosome collision. (A) Schematic representation of disome-free areas surrounding start codons. (B) Gene ontology analysis conducted with iPAGE (Goodarzi et al., 2009).

**Figure S3.**
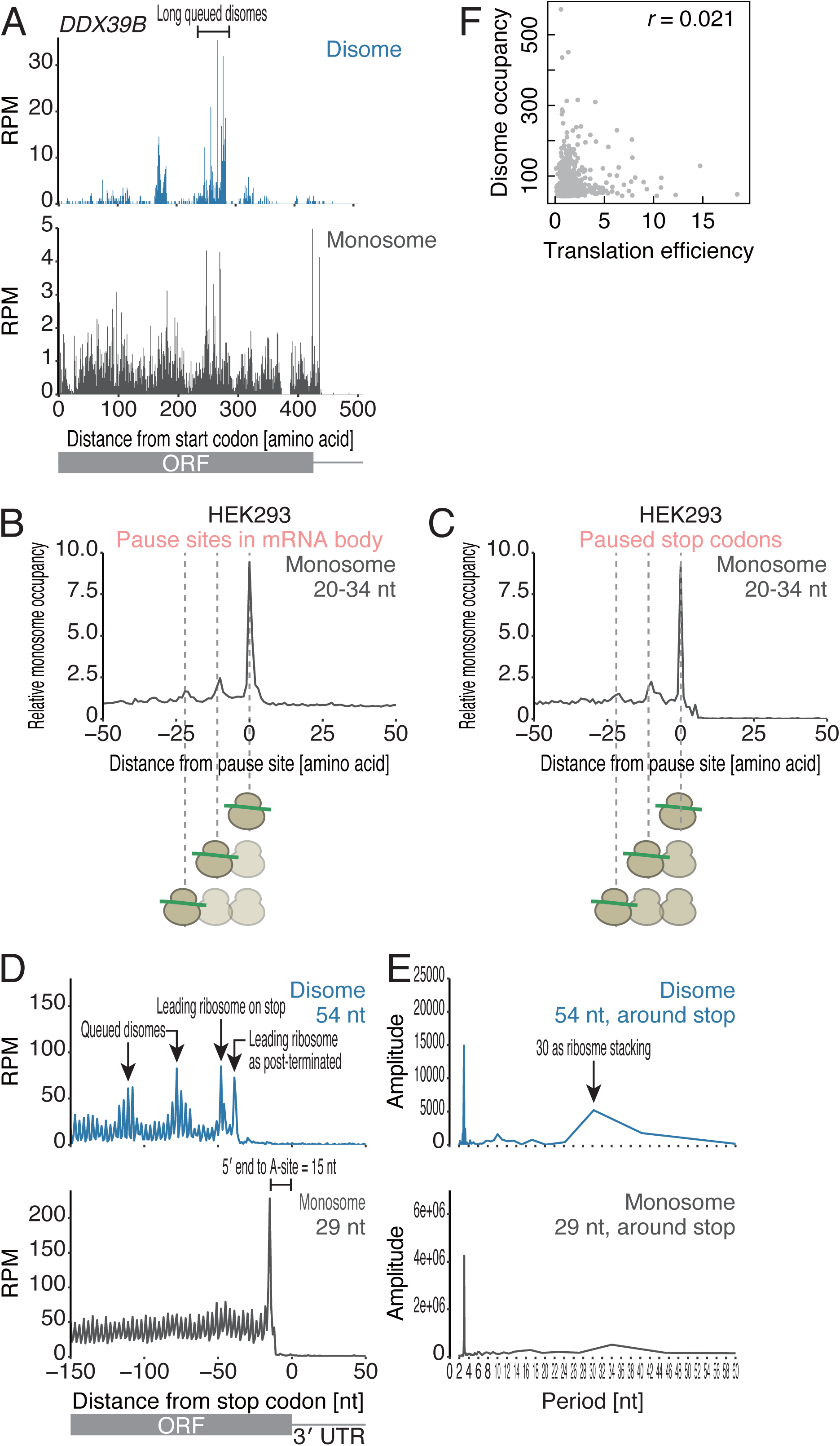
Stacking of ribosomes in long queues revealed by disome profiling. (A) The read distribution along human *DDX39B* mRNA is depicted, showing the A-site position of a monosome (bottom) and that in the leading ribosome of a disome (top). (B and C) Metagene analysis of monosome reads around ribosome collision sites in mRNA bodies (B) and on stop codons (C). (D) Metagene analysis showing the 5′ ends of disome (top) and monosome (bottom) footprint reads with indicated read lengths around the stop codon. (E) Discrete Fourier transform of reads showing their periodicity around stop codons. (F) The correlation between disome occupancy and translation efficiency.

**Figure S4.**
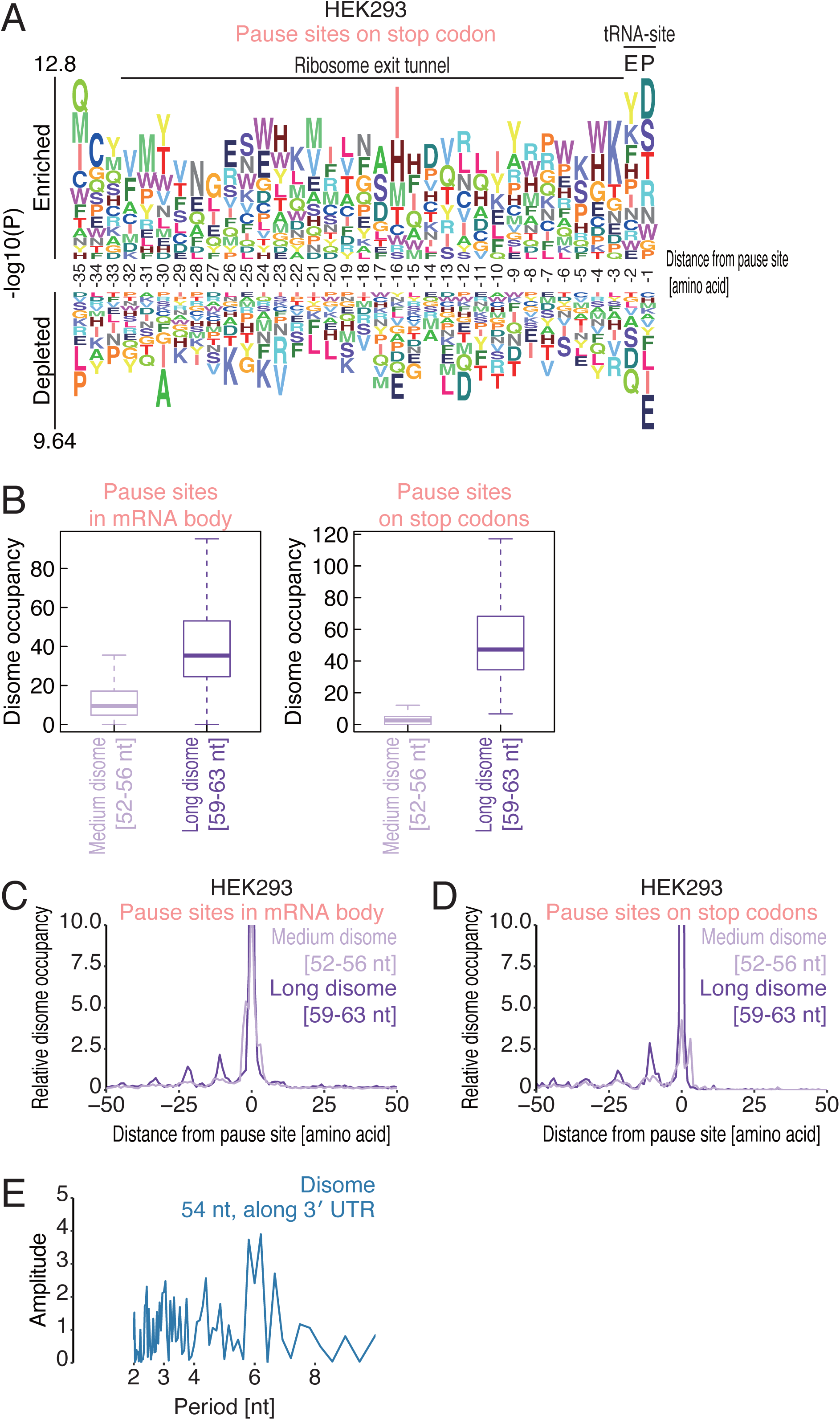
Characterization of the lengths of disome footprint reads on pause sites. (A) Amino acid sequence relative to the A-site position of the leading ribosome in a disome on a stop codon is shown. Logos were drawn by kpLogo. (B and C) Metagene analysis of medium-length (52-56 nt) and long (59-63 nt) disome footprint reads around ribosome collision sites in mRNA bodies (B) and on stop codons (C). (D) Box plots indicating the disome occupancy on pause sites indicating their footprint read lengths. (E) Discrete Fourier transform of disome footprints found in 3′ UTR.

**Figure S5.**
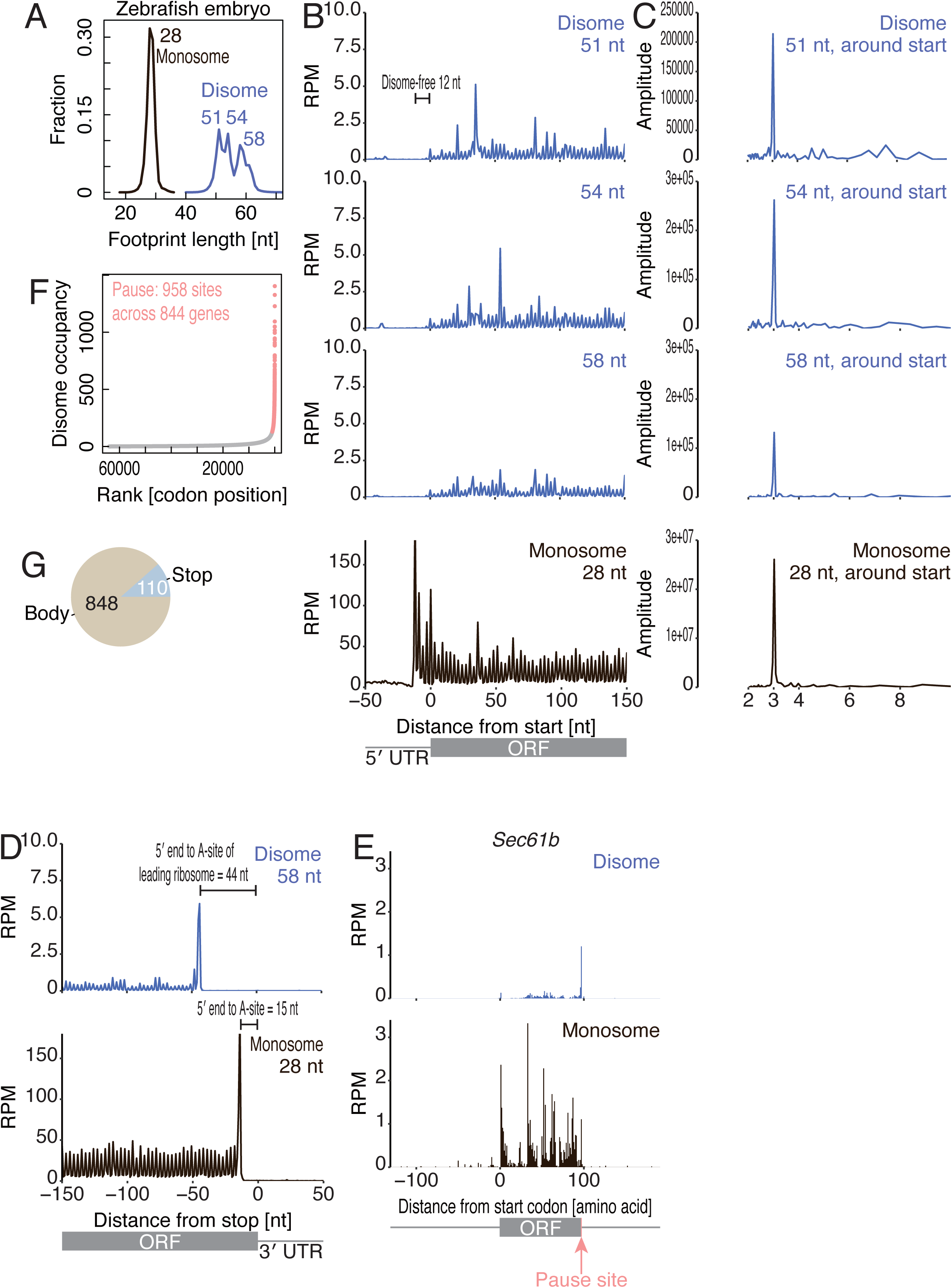
Characterization of disome profile in zebrafish. (A) Distribution of footprint read lengths generated by monosome profiling (regular ribosome profiling) and disome profiling in zebrafish embryos. (B and D) Metagene analysis showing the 5′ ends of footprints indicating read lengths around start codons (B) and stop codons (D). (C) Discrete Fourier transform of reads showing their periodicity around start codons. (E) The read distribution along *Sec61b* mRNA is depicted, showing the A-site position of a monosome (bottom) and that in the leading ribosome of a disome (top). The known ribosome pause site (Mariappan et al., 2010) is highlighted in pink. (F) Calculated disome occupancies across codons. The collision sites, which are defined as codons with a disome occupancy larger than the mean + 3 s.d., are highlighted in pink. (G) Pie chart indicating the positions of disome pause sites in mRNAs.

**Figure S6.**
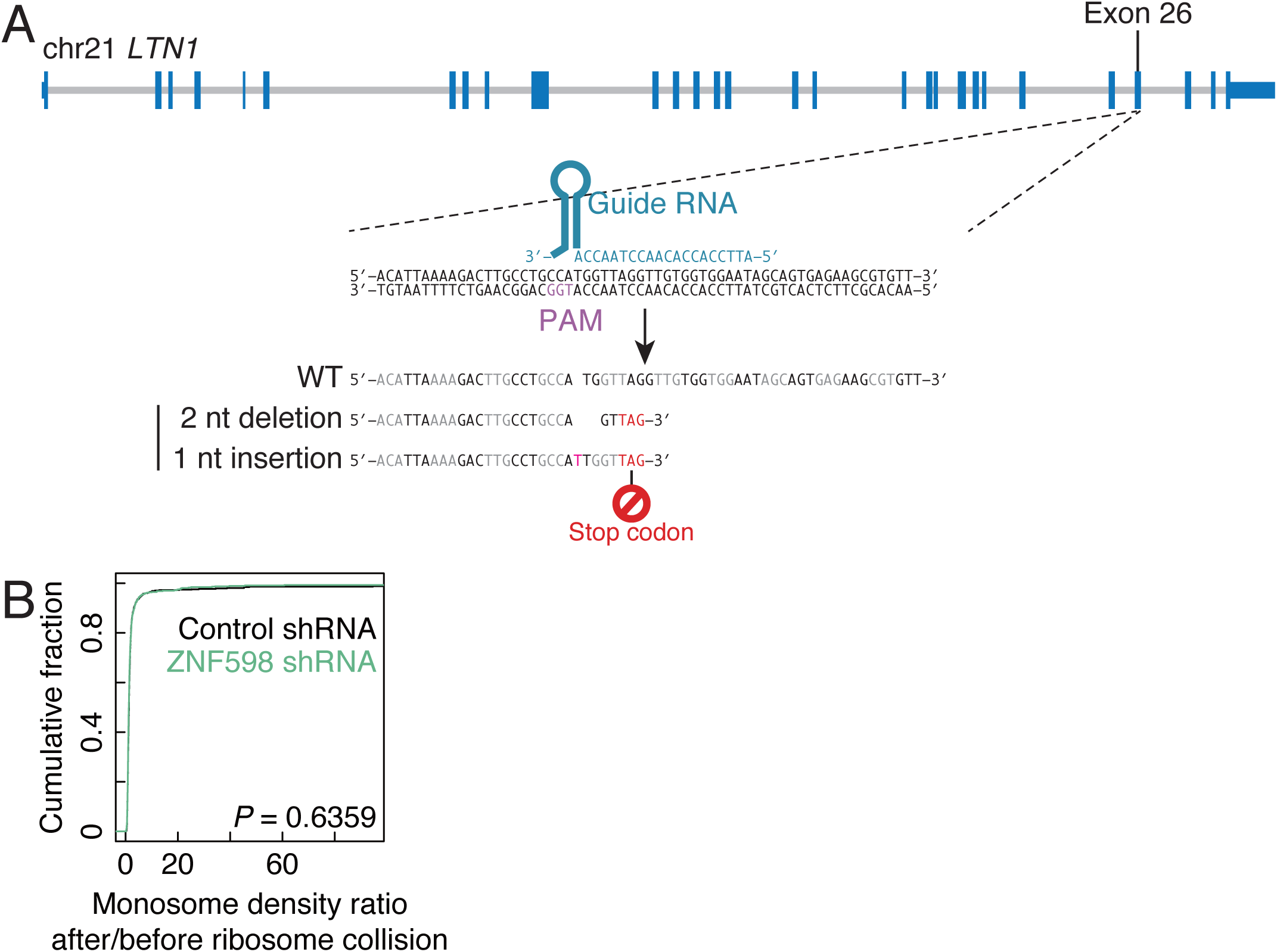
Characterization of LTN1 knockout cells and ZNF598 knockdown cells. (A) Schematic representation of gRNA designed for gene knockout and mutations by CRISPR-Cas9. (B) Cumulative distribution of the density for the monosome after the ribosome collision site to that for monosome before the ribosome collision site with or without ZNF598 knockdown.

**Figure S7.**
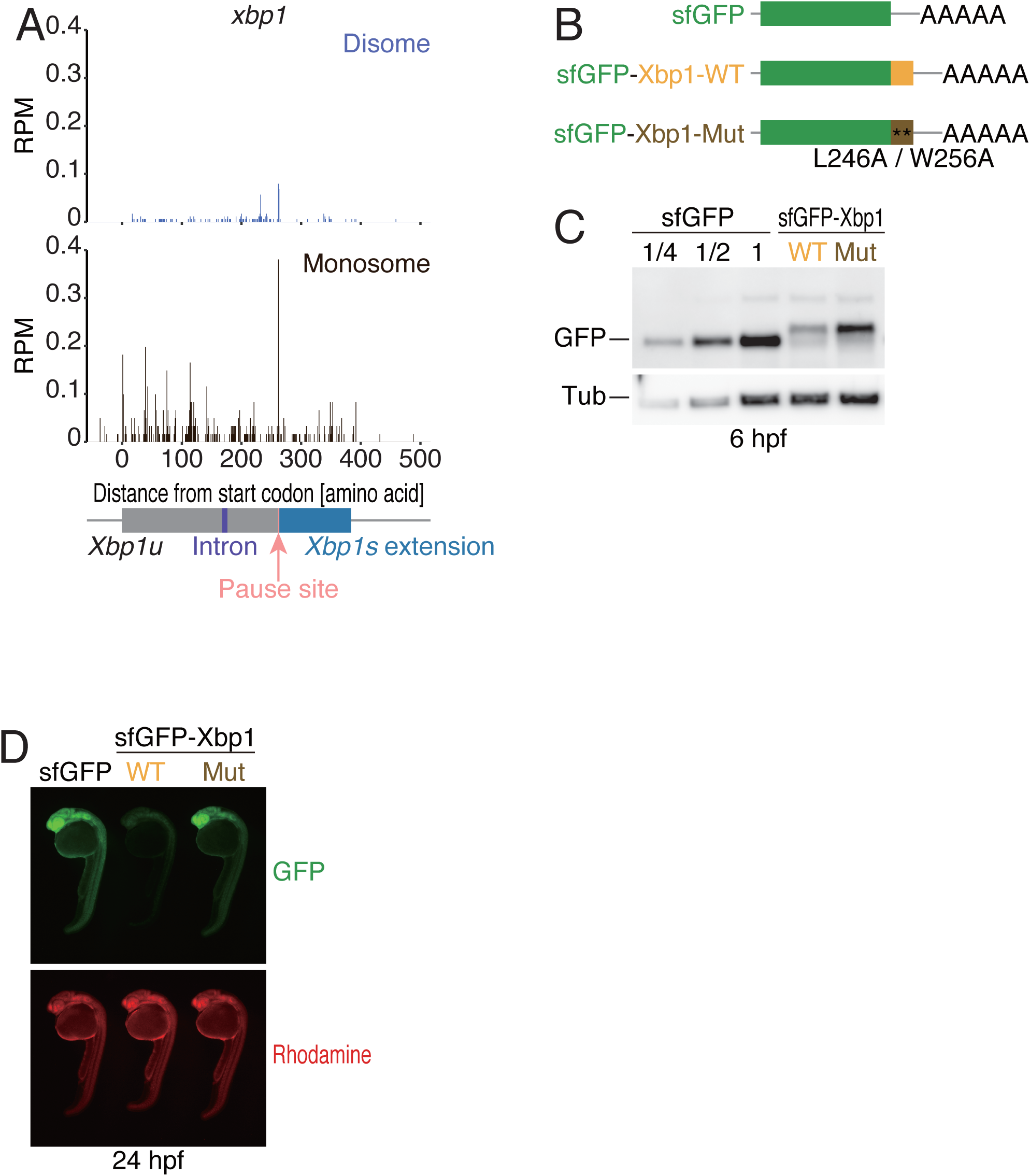
Characterization of the zebrafish *xbp1u* pause site. (A) The read distribution along zebrafish *xbp1u* mRNA is depicted, showing the A-site position of a monosome (bottom) and that in the leading ribosome of a disome (top). The known ribosome pause site (Yanagitani et al., 2011) is highlighted in pink. (B) Schematic illustrations of mRNA reporters for embryo injection experiments. The two known residues critical for *xbp1u* pausing were substituted to alanine (Yanagitani et al., 2011). (C) Western blot for the indicated proteins from reporter-injected zebrafish embryos. hpf: hours post fertilization. (D) Microscopic visualization of GFP reporter expression (green). Rhodamine-dextran was co-injected as control (red).

**Table S1. List of determined ribosome collision sites in HEK293 cells.** Each gene is listed with its UCSC identifier, codon position (ATG as 0), disome pause score, and E-, P-, and A-site codon sequences.

**Table S2. List of determined ribosome collision sites in zebrafish embryos.** Each gene is listed with its Ensembl identifier, codon position (ATG as 0), disome pause score, and E-, P-, and A-site codon sequences.

